# *In vitro* transcribed guide RNAs trigger an innate immune response via the RIG-I pathway

**DOI:** 10.1101/275669

**Authors:** Beeke Wienert, Jiyung Shin, Elena Zelin, Kathleen Pestal, Jacob E. Corn

## Abstract

CRISPR-Cas9 genome editing is revolutionizing fundamental research and has great potential for the treatment of many diseases. While editing of immortalized cell lines has become relatively easy, editing of therapeutically relevant primary cells and tissues can remain challenging. One recent advancement is the delivery of a Cas9 protein and an *in vitro* transcribed (IVT) guide RNA (gRNA) as a precomplexed ribonucleoprotein (RNP). This approach allows editing of primary cells such as T cells and hematopoietic stem cells, but the consequences beyond genome editing of introducing foreign Cas9 RNPs into mammalian cells are not fully understood. Here we show that the IVT gRNAs commonly used by many laboratories for RNP editing trigger a potent innate immune response that can be several thousand times stronger than benchmark immune stimulating ligands. IVT gRNAs are recognized in the cytosol through the RIG-I pathway but not the MDA5 pathway, thereby triggering a type I interferon response. Removal of the 5’-triphosphate from gRNAs ameliorates inflammatory signaling and prevents the loss of viability associated with genome editing in hematopoietic stem cells. The potential for Cas9 RNP editing to induce a potent antiviral response indicates that care must be taken when designing therapeutic strategies to edit primary cells.

**Abbreviations:**

Cas: CRISPR-associated
CIP: calf intestinal alkaline phosphatase
CRISPR: clustered, regularly interspaced, short palindromic repeat
dCas9: nuclease-dead Cas9
HEK293: Human embryonic kidney cells 293
HEK293T: Human embryonic kidney cells 293 SV40 large T antigen
HeLa: Henrietta Lacks cells
HSPCs: CD34^+^ human hematopoietic stem and progenitor cells
IFNAR1: Interferon Alpha And Beta Receptor Subunit 1
IFNβ/IFNB1: Interferon beta
ISG15: Interferon-stimulated gene 15 IVT – *in vitro* transcribed
KO: knockout
MAVS: mitochondrial activator of virus signaling
MDA5/IFIH1: melanoma differentiation-associated gene 5/ Interferon Induced with Helicase C Domain 1
PAMP: pathogen-associated molecular pattern
RIG-I/DDX58: retinoic acid-inducible gene I/ DExD-H-box helicase 58
gRNA: guide RNA
SPRI: solid phase reversible immobilization
WT: wild type

## Introduction

CRISPR (clustered, regularly interspaced, short palindromic repeat)-Cas (CRISPR-associated) genome editing has rapidly become a widely used tool in molecular biology laboratories. Its ease of use and high flexibility allows researchers to modify and edit genomes in cell lines (1), stem cells (2), animals and plants (3,4), and even human embryos (5). At least two components must be successfully delivered into cells during genome editing: the Cas protein, such as Cas9, and a guide RNA (gRNA) to direct the Cas protein to its target site. For *in vitro* cultured cells this can be done by transfecting plasmids encoding gRNA and Cas9 protein. However, transfection of plasmid DNA into sensitive cell types such as primary and stem cells is challenging and inefficient. The introduction of plasmids can also lead to undesired integration of DNA at the cut site (6), increased off-target activity through prolonged expression of the CRISPR-Cas9 components (7), and a delay in editing while the cell expresses gRNA and Cas protein (8).

The delivery of gRNA and Cas9 protein as a pre-complexed ribonucleoprotein (RNP) sidesteps issues related to plasmid expression and has proved to be a successful strategy to edit human primary cells, including T cells (9,10), hematopoietic stem cells (11–14), and neurons (15). This makes RNP editing a particularly attractive approach for therapeutic applications, but relatively little is known about the non-editing consequences of introducing a foreign gRNA and Cas9 protein. Human cells have evolved multiple defense mechanisms to guard against foreign components, and genome editing reagents have the potential to activate these systems. For example, recent data suggests that humans may have a pre-existing adaptive immune response to the Cas9 protein (16). But cellular responses to the gRNAs used to program Cas9 editing have so far not been well explored.

Cells respond to infection by RNA-viruses with an innate immune response that protects the host cell from invading foreign genetic material (17). Foreign RNAs are recognized by pathogen-associated molecular pattern (PAMP) binding receptors in the cytosol that include retinoic acid-inducible gene I (RIG-I) and melanoma differentiation-associated gene 5 (MDA5) (18). This triggers a cascade of events mediated by the mitochondrial activator of virus signaling (MAVS) protein resulting in the transcriptional activation of type I interferons and interferon-stimulated genes (ISGs) (19–21).

PAMPs usually contain exposed 5’-triphosphate ends (18), which may also be present in gRNAs made via T7 *in vitro* transcription (IVT) (22,23). We asked whether IVT gRNAs cause an innate immune response, and here show that RNP genome editing induces upregulation of interferon beta (IFNβ) and interferon-stimulated gene 15 (ISG15) in a variety of human cell types. This activity depends upon RIG-I and MAVS, but is independent of MDA5. The extent of the immune response depends upon the protospacer sequence, but removal of the 5’-triphosphate from gRNAs avoids stimulation of innate immune signaling. The potential for Cas9 RNP editing to induce an antiviral response indicates that care must be taken when designing therapeutic strategies to edit primary cells.

## Results

To investigate if mammalian cells react to IVT gRNA/Cas9 with an innate immune response, we first performed genome editing in human embryonic kidney 293 (HEK293) cells using Cas9 RNPs. To separate innate immune response from genome editing, we performed these experiments with a non-targeting gRNA that recognizes a sequence within BFP and has no known targets within the human genome (24). Constant amounts of recombinant Cas9 protein were complexed with different amounts of non-targeting IVT gRNA and RNPs were transfected into HEK293 cells using CRISPRMAX lipofection reagent (25). We harvested cells 30h after transfection and measured transcript levels of *IFNB1* and *ISG15* by qRT-PCR (**Figure 1A**). Introduction of gRNAs caused a dramatic increase in both *IFNB1* and *ISG15* levels, and the presence of Cas9 protein did not have an effect on the outcome. Cas9 on its own did not induce *IFNB1* or *ISG15* expression. To our surprise, as little as 1 pmol of gRNA was sufficient to trigger a 30-50-fold increase in the transcription of innate immune genes. We further found that the commonly-administered 100 pmol of a gRNA can induce *IFNB1* by 4,000 fold, which is equal to or even stronger than induction by canonical IFNβ inducers such as viral mRNA from Sendai Virus (26) or a Hepatitis C virus (HCV) PAMP (20,27) (**Figure 1B**).

**Figure 1:**
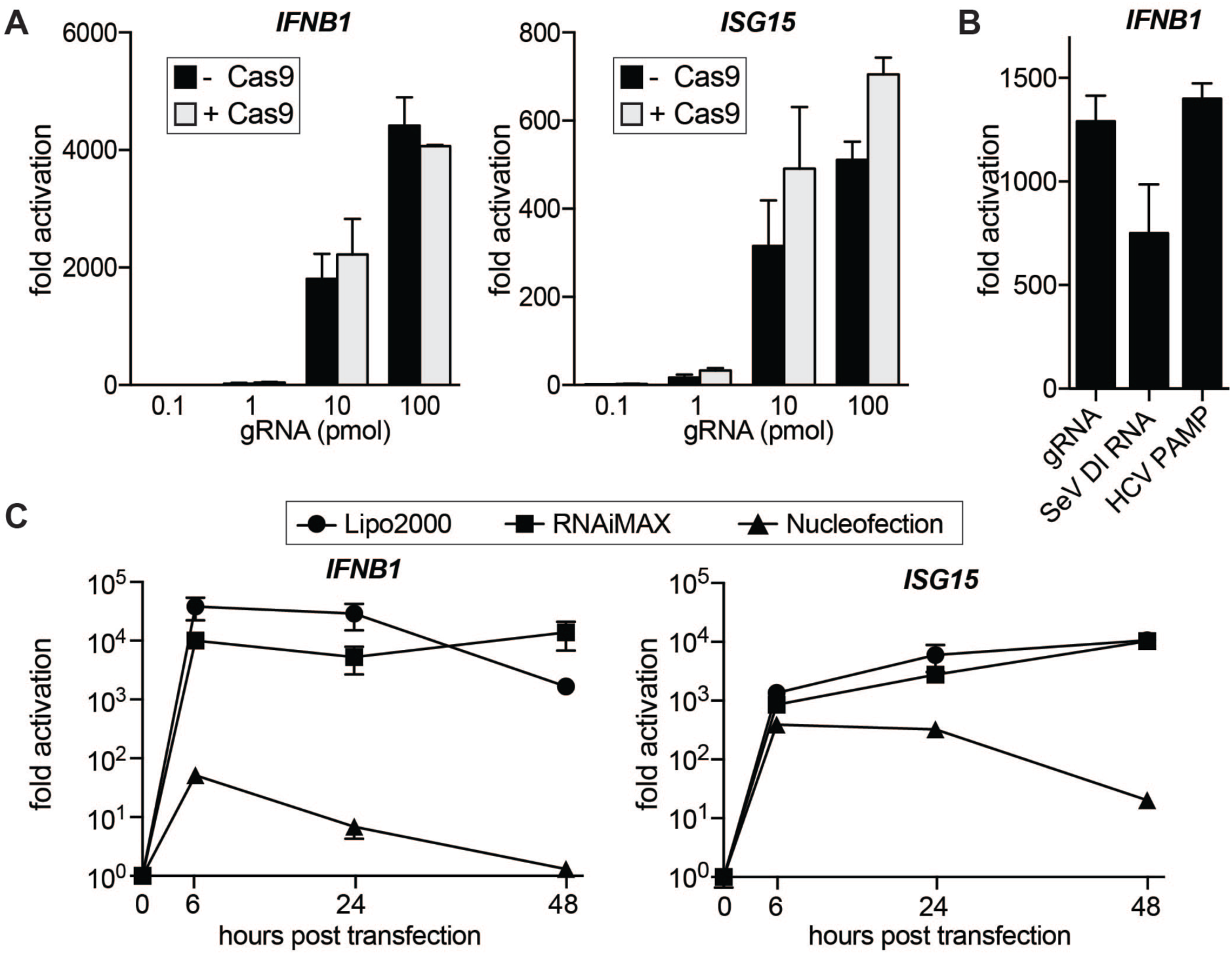
Transfection of IVT gRNAs into HEK293 cells triggers a type I interferon response. **(A)** qRT-PCR analysis of *IFNB1* and *ISG15* transcript levels in HEK293 cells transfected with increasing amounts of gRNA with and without Cas9 protein. In the samples with Cas9, gRNAs were complexed with constant amounts (100 pmol) of Cas9 protein. Cells were harvested for RNA extraction 30h after transfection using CRISPRMAX transfection reagent. Ct values were normalized to Ct values of mock transfected HEK293 cells to determine fold activation. (**B**) qRT-PCR analysis of *IFNB1* transcript levels in HEK293 cells transfected with equimolar amounts (50 pmol) of IVT gRNA, Sendai Virus defective interfering (SeV DI) RNA or HCV PAMP,respectively. (**C**) qRT-PCR analysis of *IFNB1* and *ISG15* transcript levels in HEK293 cells over a 48h time course after transfection with 50 pmol via lipofection (Lipofectamine2000 or RNAiMAX) or nucleofection, respectively. For all panels average values of three biological replicates +/-SD are shown.

RNPs can be delivered into cells via different transfection methods and while lipofection is cost-effective and easy to use, many researchers prefer electroporation for harder-to-transfect cells. We wondered if the transfection method would affect the IFNβ response and compared gRNA transfection via lipofection (Lipofectamine 2000 and RNAiMAX) to nucleofection (Lonza) (**Figure 1C**). Lipofection led to a strong increase in *IFNB1* and *ISG15* transcript levels after as little as 6h post transfection, and the response was sustained for up to 48 hours. Nucleofection also caused an increase in innate immune signaling at early time points, but the response was much milder than in lipofected samples and was greatly diminished by 48 hours.

Next, we asked if the innate immune response to gRNAs is a common phenomenon across different cell types and compared IFNβ activation in seven commonly used human cell lines of various lineages: HEK293T, HEK293, HeLa, Jurkat, HCT116, HepG2 and K562 (**Figure 2A**). While the magnitude of induction varied between cell lines, all tested cell lines responded to IVT gRNA transfection with activation of *IFNB1* expression. The sole exception was K562 cells, which have a homozygous deletion of the *IFNA* and *IFNB1* genes (28). We also measured transcript levels of two major cytosolic pathogen recognition receptors, RIG-I (*DDX58*) and MDA5 (*IFIH1*), and noticed that all cell lines except K562 upregulated these transcripts in response to introduction of gRNAs. We also confirmed these results on the protein level in HEK293 cells (**Figure 2B**).

**Figure 2:**
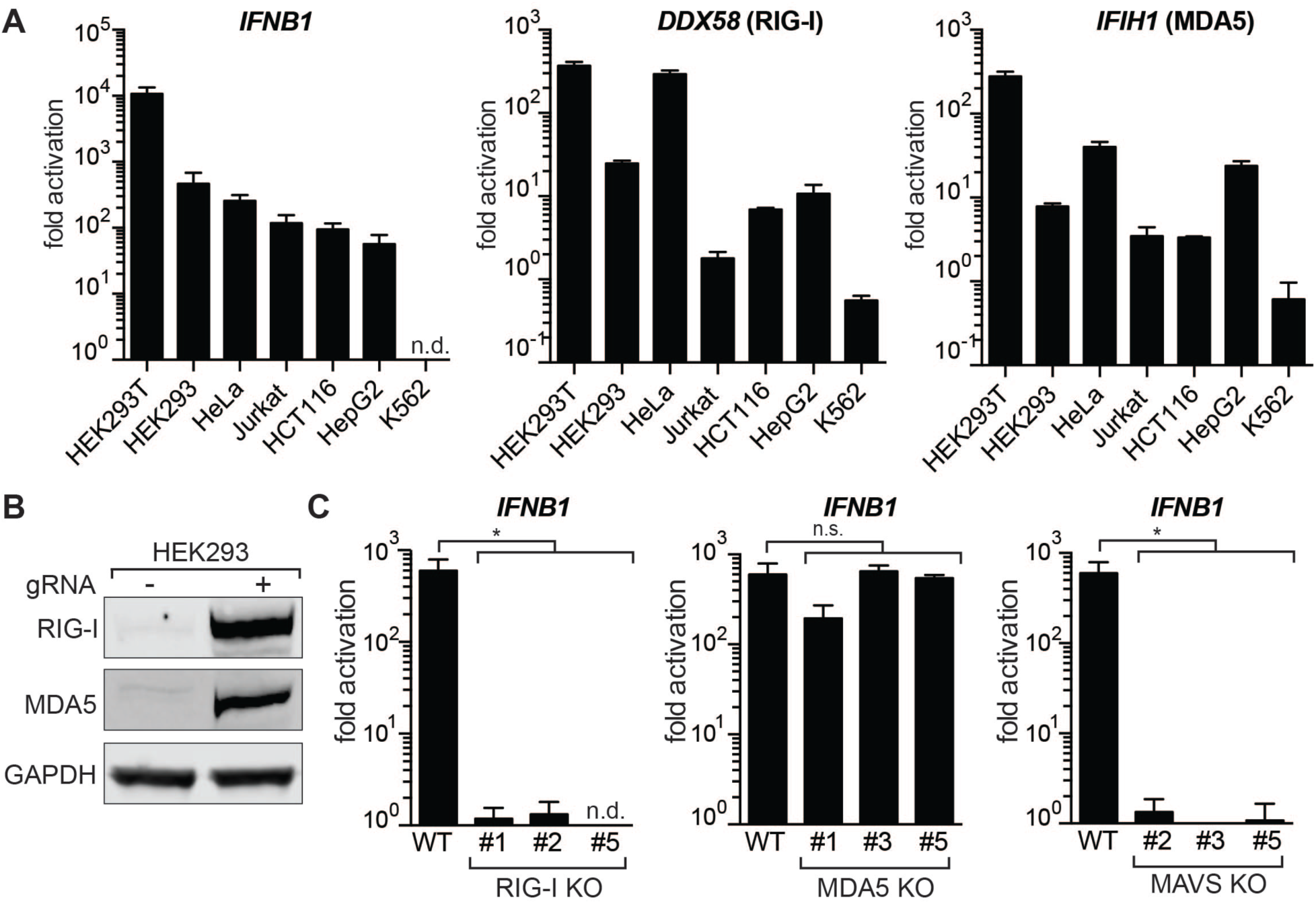
IVT gRNAs are recognized via the RIG-I pathway. (**A**) qRT-PCR analysis of increase in *IFNB1* transcript levels (left) and transcript levels of the two main cytosolic RIG-I-like receptors (*DDX58* and *IFIH1*) after introduction of IVT gRNA. Cell lines were ordered by responsiveness to gRNA-mediated induction of *IFNB1* transcript levels. Cells were harvested for RNA extraction 30h after transfection. Ct values were normalized to Ct values of mock transfected cells for each cell line to determine fold activation. *IFNB1* levels for K562 cells were too low to be determined (n.d.). (**B**) Western Blot analysis for RIG-I and MDA5 expression of mock transfected and gRNA transfected HEK293 cells after 48h. (**C**) qRT-PCR analysis of *IFNB1* transcript levels in HEK293 RIG-I (left), MDA5 (middle) and MAVS (right) knockout cells. Shown are three biological replicates of three clonal populations of RIG-I, MDA5 or MAVS KO cells, respectively. *IFNB1* levels for RIG-I KO clone #5 were too low to be determined (n.d.). For panels A and C, cells were harvested for RNA extraction 30h after transfection using RNAiMAX transfection reagent. Average values of three biological replicates +/-SD are shown. Statistical significances were calculated by unpaired t-test (*p<0.0001, n.s.: not significant).

The RIG-I and MDA5 receptors complement each other by recognizing different structures in foreign cytosolic RNAs, but the exact nature of their ligands is not yet fully understood (29,30). To investigate if IVT gRNAs are recognized via RIG-I or MDA5, we generated clonal knockout (KO) cell lines for RIG-I, MDA5, and their downstream interaction partner MAVS in HEK293 cells using CRISPR-Cas9. As the expression of both RIG-I and MDA5 are themselves stimulated by IFNβ, we confirmed successful KO after transfection with gRNAs by genomic PCR, Sanger sequencing, and Western Blot (**Supplementary Figure 1A-C**). MAVS KO cells were confirmed by Western Blot (**Supplementary Figure 1D**). Strikingly, activation of *IFNB1 e*xpression after introduction of gRNAs was absent in RIG-I and MAVS KO cells, while MDA5 KO cells did not show a significant decrease in *IFNB1* transcript levels (**Figure 2C**). This indicates that IVT gRNAs are exclusively recognized through RIG-I to trigger a type I interferon response.

As the structural requirements of RIG-I ligands are still not completely understood, we wondered if different 20 nucleotide protospacers in gRNAs vary in their potency to trigger an innate immune response via RIG-I. We designed 10 additional non-targeting gRNAs that we *in vitro* transcribed and transfected into HEK293 cells. Surprisingly, we found that the cells responded to different protospacers with a wide range of differential *IFNB1* expression. Several gRNAs produced very little innate immune response, and one gRNA (gRNA11) yielded no *IFNB1* activation at all (**Figure 3A**). We speculated that the differential response may be correlated with the purity of the RNA product after IVT or the stability of the secondary structure of the RNA (31). However, we found that there was no obvious correlation between the immune response to certain gRNAs and their purity, predicted protospacer secondary structure, full secondary structure including the constant region, or predicted disruption of the constant region by mis-pairing with the protospacer (**Supplementary Figure 2**). These results indicate that RIG-I recognition patterns of IVT gRNAs are complex and difficult to predict *a priori* based on predicted properties of the variable protospacer.

**Figure 3.**
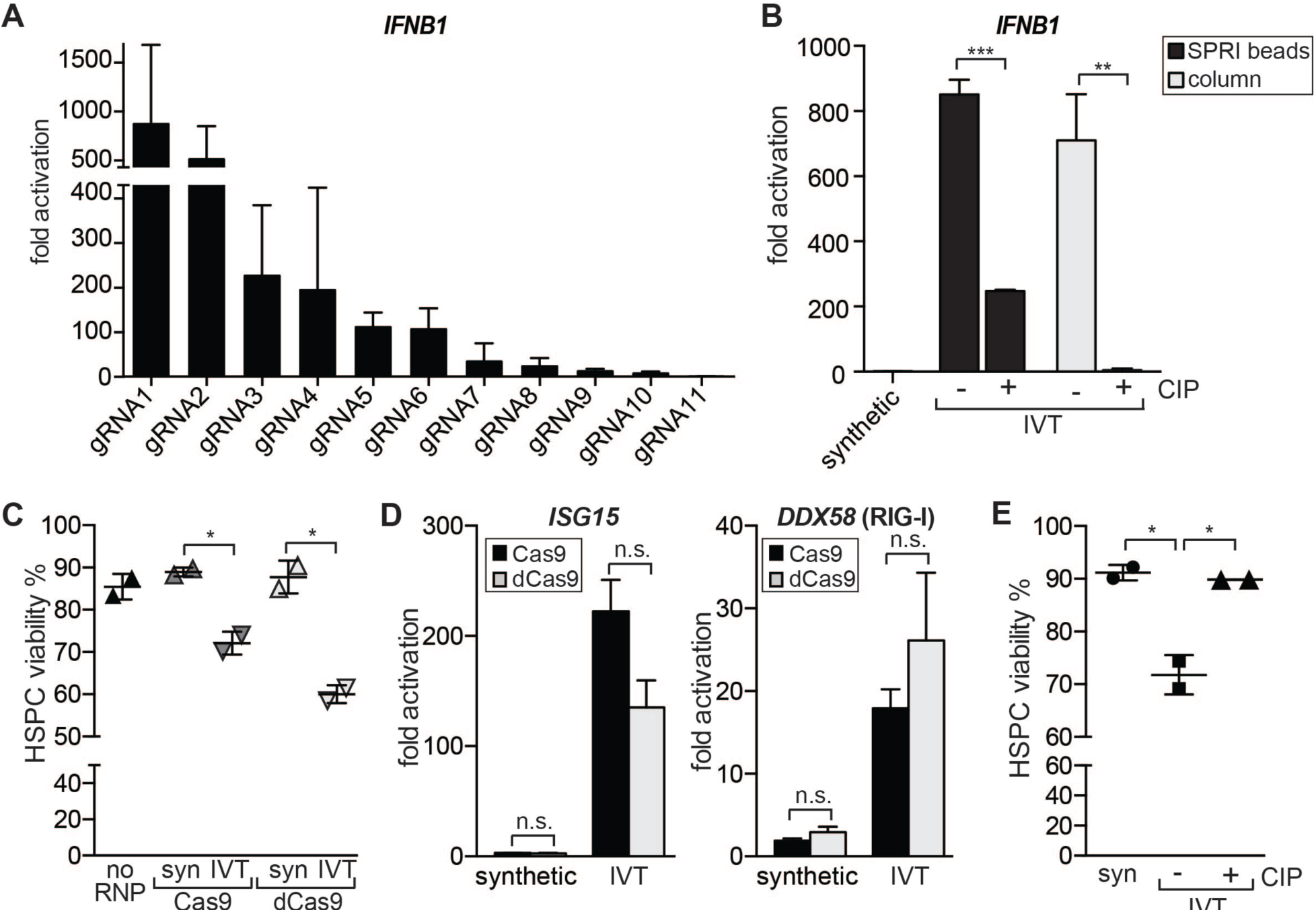
Protospacer and 5’-triphosphate determine the intensity of the gRNA-mediated IFNβ response. **(A)** qRT-PCR analysis of *IFNB1* transcript levels in HEK293 cells transfected with equal amounts of gRNAs containing different 20 nucleotide protospacers. gRNAs were ordered by decreasing levels of *IFNB1* activation. gRNA1 refers to the gRNA that has been used in all previous experiments. **(B)** qRT-PCR analysis of *IFNB1* transcript levels in HEK293 cells transfected with synthetic, IVT and Calf intestine phosphatase treated (CIP) IVT gRNAs (gRNA1). After IVT and CIP treatment gRNAs were purified with SPRI beads or spin columns, respectively. Cells were harvested for RNA extraction 30h after transfection with RNAiMAX transfection reagent. Average values of three biological replicates +/-SD are shown. **(C)** Viability of human primary HSPCs 24h post nucleofection with no RNP and Cas9 or dCas9 RNPs. dCas9 or Cas9 were complexed with synthetic (syn) or IVT gRNA targeting the *HBB* gene. Viability was determined by Trypan blue exclusion test. (**D**) qRT-PCR analysis of *ISG15* and *DDX58* (RIG-I) transcript levels in human primary HSPCs 16h post nucleofection. dCas9 or Cas9 were complexed with synthetic or IVT gRNA targeting the *HBB* gene, respectively. Ct values were normalized against Ct of mock-nucleofected cells. Average values of two biological replicates +/-SD are shown. (**E**) Viability of human primary HSPCs 16h post transfection with RNPs. RNPs consisted of dCas9 complexed with synthetic, IVT or CIP-treated IVT gRNAs targeting a non-coding intron of *JAK2*. Viability was determined by Trypan blue exclusion test. Statistical significances were calculated by unpaired t-test (*p<0.05, **p<0.01, ***p<0.0001, n.s.: not significant).

One well-established requirement of RIG-I ligands is the presence of a 5’-triphosphate group (32). We asked if preparations that remove the 5’ triphosphate might avoid or reduce the innate immune response to IVT gRNAs. We first used a synthetic gRNA that lacks a 5’-triphosphate and verified that this gRNA does not induce *IFNB1* expression when transfected into HEK293 cells (**Figure 3B**). Synthetic guide RNAs are becoming more commonplace, but are still an order of magnitude more expensive than IVT of gRNAs. This limits their application for high-throughput interrogation of gene function in primary cells. We therefore asked if treatment of IVT gRNA with calf intestinal alkaline phosphatase (CIP) to remove the 5’-triphosphate would reduce *IFNB1* induction. We found that CIP treatment significantly reduced the *IFNB1* response (**Figure 3B**). Using different CIP treatment regimes, we further found that removal of the 5’-triphosphate must to be absolutely complete and should be carried out rigorously to avoid IFN stimulation (**Supplementary Figure A-B**). We also compared purification of IVT gRNAs by solid phase reversible immobilization (SPRI) beads to column purification and established that SPRI bead clean-up is not sufficient to completely avoid an immune response even when more CIP is used (**Figure 3B and Supplementary Figure 3C**). Taken together, these results indicate that 5’-triphosphate is a necessary requirement for gRNA-induced *IFNB1* activation through RIG-I, but that additional structural properties of the gRNAs also influence the magnitude of the immune response.

When a cell initiates an antiviral immune response, it also undergoes cellular stress that can affect cell viability (33,34). Hence, we asked if there is a correlation between the IFNβ response and cell viability after transfection with synthetic, IVT or CIP-treated IVT gRNA. Not surprisingly, the viability of the very robust HEK293 cell line was not affected by the antiviral immune response (**Supplementary Figure 3D**). We then turned to primary CD34^+^ human hematopoietic stem and progenitor cells (HSPCs), which are a much more sensitive cell type. We first nucleofected HSPCs with RNPs targeting the *HBB* gene (11) and compared synthetic and IVT gRNA interferon stimulation and cell viability post transfection. Double-strand breaks have been reported to cause innate immune stimulation and can themselves cause decreases in cell fitness (35,36). Therefore, we performed controls using nuclease-dead Cas9 (dCas9) to form RNPs. We saw a significant decrease in cell viability of HSPCs in both IVT gRNA RNP samples which was associated with an increase in IFN stimulated genes *ISG15* and *RIG-I* (**Figure 3C-D**). We did not see a substantial difference in viability or ISG expression between Cas9 and dCas9 RNPs suggesting that nuclease activity did not affect the immune response. Next, we asked if CIP treatment of gRNAs could reverse the decrease in viability in HSPCs. We nucleofected HSPCs with dCas9 RNPs targeting a non-coding intron of *JAK2* and compared synthetic, IVT and CIP-treated IVT gRNAs. Strikingly, CIP treatment could completely restore the viability in HSPCs (**Figure 3E**).

## Discussion

We have found that IVT gRNAs used with Cas9 RNPs for many genome editing experiments can trigger a strong innate immune response in many mammalian cell types (**Figure 4**). Lipofection results in a stronger and longer lasting response than nucleofection, possibly because lipofection delivers gRNAs to the cytosol while nucleofection delivers mainly to the nucleus. Using isogenic knockout clones, we found the gRNA-induced response is mediated via the anti-viral RIG-I pathway and results in expression of genes that initiate an antiviral immune response. While introduction of IFN-stimulating gRNAs does not affect viability in HEK293 cells, we found that viability of primary HSPCs is negatively affected by the antiviral immune response.

**Figure 4.**
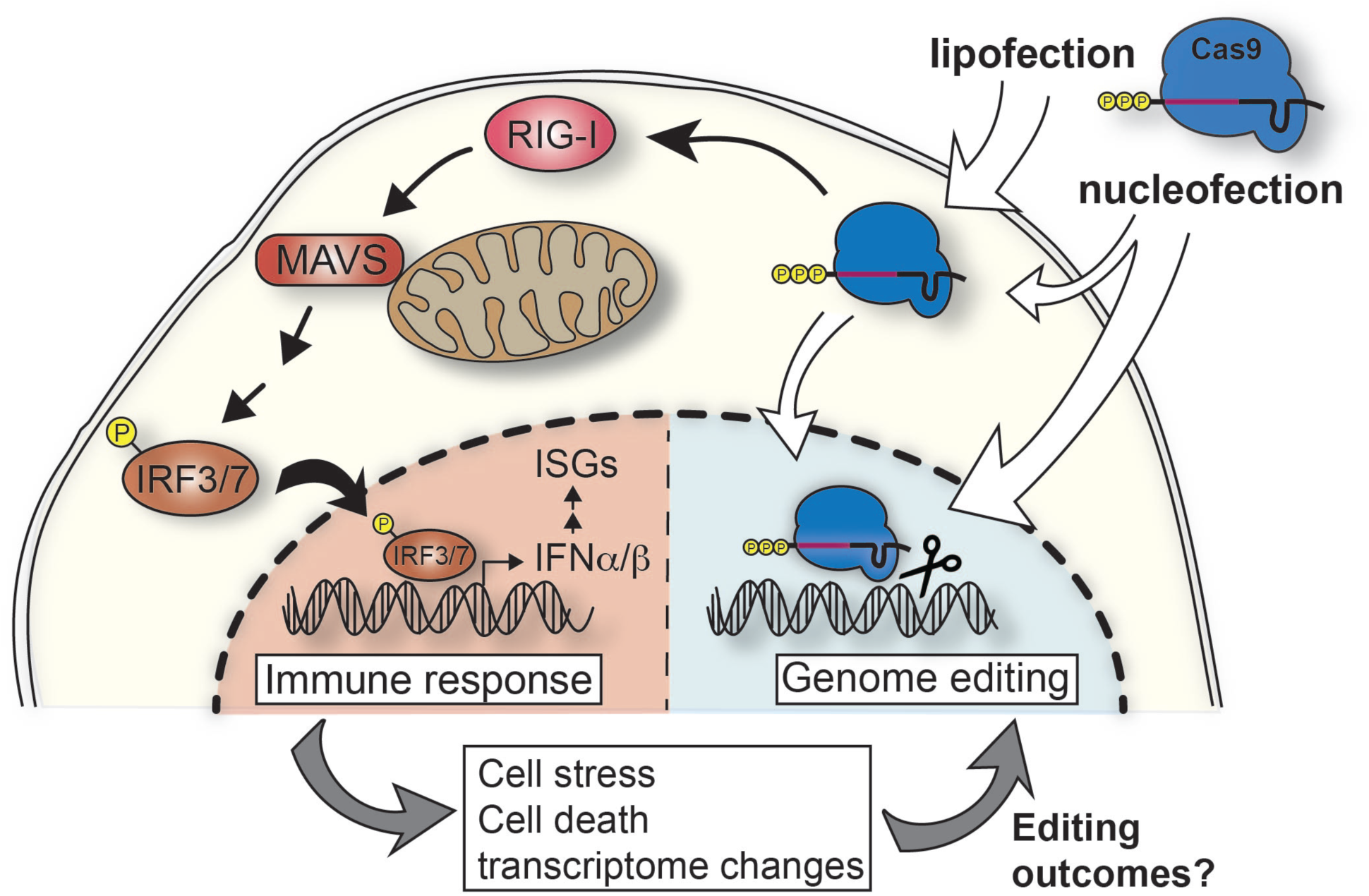
Transfection of *in vitro* transcribed gRNAs induces a cytosolic immune response. Proposed model of IVT gRNA recognition pathways in mammalian cells. IVT gRNAs carry a 5’-triphosphate and when complexed with Cas9 protein and transfected into cells, cytosolic RNPs are recognized by RIG-I triggering a cascade of activation events through the mitochondrial antiviral signaling protein (MAVS). This results in phosphorylation of IRF3/7 and their shuttling into the nucleus to activate expression of type I interferons (IFNα/β). This triggers the expression of interferon-stimulated genes (ISGs). This innate immune response changes the transcriptome of the cell and can cause cell stress and/or death which in turn might affect the editing outcomes.

These results have several implications. We suggest that the gene signature associated with type I interferon stimulation should be considered when studying the transcriptome of recently edited bulk populations of cells. Furthermore, all mammalian cells can both produce type I interferons and also respond to them through the ubiquitously expressed receptor Interferon Alpha And Beta Receptor Subunit 1 (IFNAR1) (37). Even cells that have not been successfully transfected with RNPs could sense the INFβ produced by neighboring cells and activate downstream antiviral defense mechanisms. This could be an important consideration during *in vivo* genome editing applications as RNP delivery into one set of cells could provoke a wide-spread innate immune response in the surrounding tissues.

We found that synthetic gRNAs completely circumvent the RIG-I mediated response, offering a valuable path to avoid innate immune signaling during therapeutic editing. However, synthetic gRNAs can become expensive when performing experiments that require testing or using many gRNAs. We found that a cost-effective CIP treatment to remove the 5’-triphosphate before transfection reduces the immune response and increases post-transfection viability in HSPCs. Complete removal of the 5’-triphosphate is essential to restore viability in HSPCs to the same level as observed with synthetic gRNAs. Thus, consideration of a potential innate immune stimulation prior to choice of genome editing reagents, study design, and implementation of controls is critical when performing genome editing using RNPs in mammalian cells.

Double-strand breaks have on their own been reported to induce an innate immune response (35). However, we performed controls comparing WT Cas9 to a nuclease-dead Cas9 thereby showing that the gRNA-mediated innate immune response and associated cell death in HSPCs is not caused by double-strand breaks.

While we were preparing this manuscript for submission, the Kim group reported similar results in HeLa cells and primary human CD4^+^ T cells (38). They confirmed that the type I interferon response is dependent on the presence of a 5’-triphopsphate group and that CIP treatment can increase viability by avoiding the antiviral response. These results are very much in alignment with our findings and extend the potential problem of innate immune signaling to additional cell types.

Our study adds extra depth by further outlining the mechanisms by which gRNAs are sensed. We show that gRNA sensing depends upon RIG-I and MAVS, but MDA5 knockout cells are fully capable of inducing IFNβ after IVT gRNA transfection. Hence, gRNA sensing is independent of the MDA5 PAMP receptor. Furthermore, we show that in addition to a 5’-triphosphate, the protospacer sequence is also critical to determine the intensity of the IFNβ response. Not only do different gRNAs induce different innate immune responses, some gRNAs induce no response at all. It has been proposed that 5’-basepaired RNA structures are required to activate antiviral signaling via RIG-I, but we found no correlation between signaling and a variety of predicted RNA properties, including secondary structure (31). Our results therefore suggest that the mechanism of gRNA sensing by the RIG-I pathway is relatively complex, in that it requires 5’ triphosphates but that this moiety is not sufficient to induce the response. Future work to delineate the full set of molecular features responsible for gRNA activation of innate immunity might yield accurate predictors of innate immune signaling in general.

## Materials and methods

### *In vitro* transcription of gRNAs

gRNA was synthesized by assembly PCR and *in vitro* transcription as previously described (11). Briefly, a T7 RNA polymerase substrate template was assembled by PCR from a variable 58-59 nt primer containing T7 promoter, variable gRNA guide sequence, and the first 15 nt of the non-variable region of the gRNA (T7FwdVar primers, 10 nM, Supplementary Table 1; Supplementary Table 2 for gRNA sequences), and an 83 nt primer containing the reverse complement of the invariant region of the gRNA (T7RevLong, 10 nM), along with amplification primers (T7FwdAmp, T7RevAmp, 200 nM each). The two long primers anneal in the first cycle of PCR and are then amplified in subsequent cycles. Phusion high-fidelity DNA polymerase was used for assembly (New England Biolabs). Assembled template was used without purification as a substrate for *in vitro* transcription by T7 RNA polymerase using the HiScribe T7 High Yield RNA Synthesis kit (New England Biolabs) following the manufacturer’s instructions. Resulting transcription reactions were treated with DNAse I (New England Biolabs), and RNA was purified by treatment with a 5X volume of homemade solid phase reversible immobilization (SPRI) beads (comparable to Beckman-Coulter AMPure beads) and elution in RNAse-free water.

### CIP treatment of IVT gRNAs

When gRNAs were treated with Alkaline Calf intestinal phosphatase (CIP) (New England Biolabs), 20U of CIP (2 μl) were added per 20 μl IVT reaction and samples were incubated at 37C for 3h before proceeding to purification and DNAseI treatment. CIP-treated samples and corresponding no CIP IVT gRNA controls were purified using a Qiagen RNeasy Mini Kit (Qiagen). The detailed protocol and additional notes are available online (dx.doi.org/10.17504/protocols.io.nghdbt6).

### *In vitro* transcription of HCV PAMP and Sendai Virus DI RNA

HCV PAMP IVT template (20) was generated by annealing HCV fwd and rev (5 μM each) oligos (Supplementary Table 1). 2 μl of the annealed product was used as DNA template in the subsequent IVT reaction using HiScribe T7 High Yield RNA Synthesis kit (New England Biolabs).

The plasmid containing the SeV DI RNA(26) was a gift from Prof. Peter Palese, Icahn School of Medicine at Mount Sinai, New York. Plasmid was digested with HindII/EcoRI before IVT with HiScribe T7 High Yield RNA Synthesis kit (New England Biolabs). The sequence of the IVT DI, including the T7 promoter, hepatitis delta virus ribozyme, and the T7 terminator, is:

TAATACGACTCACTATA**ACCAGACAAGAGTTTAAGAGATATGTATCCTTTTAAATTTCTTGTCTTCTTGTAAGTTTTTCTTACTATTGTCATATGGATAAGTCCAAGACTTCCAGGTACCGCGGAGCTTCGATCGTTCTGCACGATAGGGACTAATTATTACGAGCTGTCATATGGCTCGATATCACCCAGTGATCCATCATCAATCACGGTCGTGTATTCATTTTGCCTGGCCCCGAACATCTTGACTGCCCCTAAAATCTTCATCAAAATCTTTATTTCTTTGGTGAGGAATCTATACGTTATACTATGTATAATATCCTCAAACCTGTCTAATAAAGTTTTTGTGATAACCCTCAGGTTCCTGATTTCACGGGATGATAATGAAACTATTCCCAATTGAAGTCTTGCTTCAAACTTCTGGTCAGGGAATGACCCAGTTACCAATCTTGTGGACATAGATAAAGATAGTCTTGGACTTATCCATATGACAATAGTAAGAAAAACTTACAAGAAGACAAGAAAATTTAAAAGGATACATATCTCTTAAACTCTTGTCTGGT**GGCCGGCATGGTCCCAGCCTCCTCGCTGGCGCCGGCTGGGCAACATTCCGAGGGGACCGTCCCCTCGGTAATGGCGAATAGCATAACCCCTTGGGGCCTCTAAACGGGTCTTGAGGGGTTTTTTG.

The sequence of the SeV DI is highlighted in boldface.

Both, HCV PAMP and SeV DI RNA were purified by treatment with a 5X volume of homemade solid phase reversible immobilization (SPRI) beads (comparable to Beckman-Coulter AMPure beads) and elution in RNAse-free water.

### Synthetic gRNAs

Chemically synthesized gRNAs, which were purified using high-performance liquid chromatography (HPLC), were purchased from Synthego.

### RNA quality control

IVT gRNAs were analyzed using a Bioanalyzer. This was performed by the UC Berkeley Functional Genomics Laboratory (FGL) core facility. gRNAs were denatured for 5 mins at 70C before analysis on bioanalyzer.

### Cas9 protein preparation

The Cas9 construct (pMJ915) contained an N-terminal hexahistidine-maltose binding protein (His6-MBP) tag, followed by a peptide sequence containing a tobacco etch virus (TEV) protease cleavage site. The protein was expressed in E. coli strain BL21 Rosetta 2 (DE3) (EMD Biosciences), grown in TB medium at 16°C for 16h following induction with 0.5 mM IPTG. The Cas9 protein was purified by a combination of affinity, ion exchange and size exclusion chromatographic steps. Briefly, cells were lysed in 20 mM HEPES pH 7.5, 1M KCl, 10mM imidazole, 1 mM TCEP, 10% glycerol (supplemented with protease inhibitor cocktail (Roche)) in a homogenizer (Avestin). Clarified lysate was bound to Ni-NTA agarose (Qiagen). The resin was washed extensively with lysis buffer and the bound protein was eluted in 20 mM HEPES pH 7.5, 100mM KCl, 300mM imidazole,1 mM TCEP 10% glycerol. The His6-MBP affinity tag was removed by cleavage with TEV protease, while the protein was dialyzed overnight against 20 mM HEPES pH 7.5, 300 mM KCl, 1 mM TCEP, 10% glycerol. The cleaved Cas9 protein was separated from the fusion tag by purification on a 5 ml SP Sepharose HiTrap column (GE Life Sciences), eluting with a linear gradient of 100 mM – 1 M KCl. The protein was further purified by size exclusion chromatography on a Superdex 200 16/60 column in 20 mM HEPES pH 7.5, 150 mM KCl and 1 mM TCEP. Eluted protein was concentrated to 40uM, flash-frozen in liquid nitrogen and stored at -80°C.

### Culture and transfection of immortalized cell lines

Cells were obtained from ATCC and verified mycoplasma-free (Mycoalert LT-07, Lonza). HEK293, HEK293T, HCT116, HepG2 and HeLa cells were maintained in DMEM supplemented with 10 % FBS and 100 μg/mL Penicillin-Streptomycin (all Gibco). K562 and Jurkat cells were maintained in RPMI supplemented with 10% FBS and 100 μg/mL Penicillin-Streptomycin.All transfections in cell lines were performed in 12-well cell culture dishes using 2x10^5^ cells per transfection. For lipofection we used Lipofectamine^TM^ CRISPRMAX(tm)-Cas9, Lipofectamine^®^ RNAiMAX or Lipofectamine® 2000 Transfection Reagent (all Invitrogen) in reverse-transfections according to the manufacturer’s protocols. Unless stated otherwise, 2x10^5^ cells were transfected with 50 pmol of RNA and harvested 24-30h post transfection for RNA extraction.

### Culture and transfection of primary HSPCs

Human CD34+ HSPCs from mobilized peripheral blood (Allcells, Inc) were thawed and cultured in StemSpan SFEM medium (StemCell Technologies) supplemented with StemSpan CC110 cocktail (StemCell Technologies) for 48 hours before nucleofection with dCas9 RNP (50pmol of dCas9, 50 pmol of gRNA). 1.5x10^5^ HSPCs were pelleted (100 x g, 10 min) and resuspended in 20 μl Lonza P3 solution, mixed with 10ul dCas9 RNP, and nucleofected using ER100 protocol in Lonza 4D nucleofector. Viability of the cells was measured 24 hours post nucleofection using Trypan blue exclusion test. RNA was harvested 16 hours post nucleofection.

### RNA extraction, cDNA synthesis and qRT-PCR

Cell cultures were washed with PBS prior to RNA extraction. Total RNA was extracted using RNeasy Miniprep columns (Qiagen) according to the manufacturer’s instructions including the on-column DNAseI treatment (Qiagen). 1 μg of total RNA was used for subsequent cDNA synthesis using iScript^TM^ Reverse Transcription Supermix (Biorad). For qRT-PCR reactions, a total of 20 ng of cDNA was used as a template and combined with primers (see Supplementary Table 3) and SsoFast^TM^ EvaGreen® Supermix (Biorad) and amplicons were generated using standard PCR amplification protocols for 40 cycles on a StepOnePlus Real-Time PCR system (Applied Biosystems). Ct values for each target gene were normalized against Ct values obtained for *GAPDH* to account for differences in loading (ΔCt). To determine ‘fold activation’ of genes ΔCt values for target genes were then normalized against ΔCt values for the same target gene for mock-treated cells (ΔΔCt).

### Generation of knockout cell lines

For CRISPR/Cas9 genome editing we used a plasmid encoding both the Cas9 protein and the gRNA. pSpCas9(BB)-2A-GFP (px458) was a gift from Feng Zhang (Addgene plasmid #48138). We designed gRNA sequences using the free CRISPR knockout design online tool from Synthego. Two different gRNA sequences were designed for RIG-I and MDA5, respectively (see Supplementary Table 3).

2x10^5^ HEK293 cells were nucleofected with 2 μg of px458 plasmids containing both targeting gRNAs in a 1:1 ratio using a Lonza 4D nucleofector (Lonza) with the manufacturer’s recommended settings. After 48h cells were harvested and subjected to fluorescence-activated cell sorting (FACS). Cells expressing high levels of GFP were single-cell sorted into 96-well plates to establish clonal populations.

For the screening process, genomic DNA from clonal populations was extracted using QuickExtract solution (Lucigen). For KO of RIG-I and MDA5 we screened clones by genomic PCR looking for a PCR product that is significantly smaller in size than that of WT HEK293 cells (see Supplementary Table 4 for primers). PCR products were then Sanger sequenced by the UC Berkeley DNA Sequencing facility using the forward primers of the PCR reaction as sequencing primers.

### Western Blot

Cells were harvested and washed with PBS. Cells were lysed in 1xRIPA buffer (EMD Millipore) for 10 mins on ice. Samples were spun down at 14000xg for 15 mins and protein lysates were transferred to a new tube. 50 μg of total protein was separated for size by SDS-PAGE and transferred to a nitrocellulose membrane. Blots were blocked in 4% skim milk in 50 mm Tris-HCl (pH 7.4), 150 mm NaCl, and 0.05% Tween 20 (TBST) and then probed for RIG-I, MDA5, MAVS or GAPDH protein using antibodies against RIG-I (D14G6), MDA5 (D74E4), MAVS (D5A9E) or GAPDH (14C10), respectively (all Cell Signaling Technologies). This was followed by incubation with secondary antibody IRDye® 800CW Donkey anti-Rabbit IgG (Li-Cor). Rainbow^TM^ protein standards (GE Healthcare) were loaded in each gel for size estimation. Blots were visualized using a Li-Cor Odyssey Clx (Li-Cor).

## Acknowledgements

We would like to thank Prof. Peter Palese from the Icahn School of Medicine at Mount Sinai, New York, for the Sendai virus DI IVT template DNA. The CRISPR/Cas9 genome-editing plasmid px458 was a gift from Feng Zhang, Broad Institute, Cambridge, MA (Addgene plasmid # 48138).

